# Innovation in NLR and TLR sensing drives the MHC-II free Atlantic cod immune system

**DOI:** 10.1101/2020.08.07.241067

**Authors:** Xingkun Jin, Bernat Morro, Ole K. Tørresen, Visila Moiche, Monica H. Solbakken, Kjetill S. Jakobsen, Sissel Jentoft, Simon MacKenzie

## Abstract

The genome sequencing of Atlantic cod revealed an immune system absent of specific cell surface toll-like receptors (TLRs), major histocompatibility complex (MHC) class II, invariant chain (CD74) and the CD4 (cluster of differentiation 4) receptor. Despite the loss of these major components considered as critical to vertebrate innate and adaptive immune systems the cod system is fully functional, however the underlying mechanisms of the immune response in cod remain largely unknown. In this study, *ex vivo* cod macrophages were challenged with various bacterial and viral microbe-associated molecular patterns (MAMP) to identify major response pathways. Cytosolic MAMP-PRR pathways based upon the NOD-like receptors (NLRs) and RIG-I-like receptors (RLRs) were identified as the critical response pathways. Our analyses suggest that internalization of exogenous ligands through scavenger receptors drives both pathways activating transcription factors like NF-kB (Nuclear factor-kappa B) and interferon regulatory factors (IRFs). Further, ligand-dependent differential expression of a unique TLR25 isoform and multiple NLR paralogues suggests (sub)neofunctionalisation toward specific immune defensive strategies. Our results further demonstrate that the unique immune system of the Atlantic cod provides an unprecedented opportunity to explore the evolutionary history of PRR-based signalling in vertebrate immunity.

## Introduction

In vertebrates, the genetic basis of immunity is considered highly conserved(1). With the increased use of high-throughput sequencing technologies, genomic resources from non-model species has become readily available and large efforts in comparative immunology has ensued (2–4). These studies have revealed several common denominators of vertebrate immunity, but also demonstrated considerable gene losses and expansions challenging our understanding of immune system organization and functional compartmentalization in vertebrates. Here, the genome sequencing of the Atlantic cod (5), and later a set of ~30 codfishes (2), demonstrated that this group of teleosts (the Gadiformes) display a very distinct immune gene repertoire affecting both innate and adaptive immunity, with the lack major histocompatibility complex (MHC) class II, invariant chain (CD74) and CD4. All hallmarks of the perceived classical vertebrate adaptive immune system (2, 5). In parallel, teleosts display a wide range of gene expansions/contractions related to major histocompatibility complex class I as well as innate immune gene families such as pattern recognition receptors (PRRs) (6–8). PRRs are a strongly conserved feature of the immune system indispensable from insects to mammals and in plants (R protein) (1, 9). Classically, these receptors are associated with detecting microbe-associated or danger-associated molecular patterns (MAMPS and DAMPS) with subsequent initiation of inflammation. However, they have also been implicated in regulation of development, antigen presentation and autophagy(10, 11).

Genomic investigations have provided detailed characterisations of all known PRR families including Toll-like receptors (TLRs), C-type lectin receptors (CLRs), retinoic acid-inducible (RIG)-I-like receptors (RLRs) and nucleotide-binding oligomerization (NOD)-like receptors (NLRs) in an array of vertebrates (9). The PRR repertoire in teleost fish has proven to be very diverse compared to other vertebrate groups, and more so within the codfishes (12, 13). Here, all homologs of mammalian surface-located TLRs have been lost, whereas there are large expansions of teleost-specific TLRs with unknown cellular location and ligand type. Furthermore, loss of RIG-I and NOD2 has been reported in parallel to the overall NLR repertoire being greatly expanded (5, 7, 12–14). These significant gene losses and expansions result in a peculiar genetic basis for innate immunity that suggests the existence of an alternative MAMP-PRR activation system. The functional implication of the PRR repertoire observed in codfishes is poorly understood. Furthermore, there is a lack of functional investigations targeting PRR interaction and signalling. So far, complete insight into these gene losses and expansions, and the expression pattern of the various paralogs, have been hampered by a fragmented genome assembly and the use of short-read technology in e.g. RNA-Seq analyses.

Certain PRR-encoding gene family groups such as the NLRs appear to contain species-specific expansions that present multiple modified forms (neo/sub-functionalization) and different genomic organizational patterns (tandem/dispersed). Atlantic cod and other gadiform species contain a high number of tandem-repeated gene families that cause assembly collapse, as a consequence PRR genes are often fragmented or collapsed (7, 12, 15). Additionally, some of the codfishes PRRs demonstrate significant gene expansions and neofunctionalization (3, 16). In the Atlantic cod genome, genes from both the TLR and NLR families were found to be highly expanded, accompanied by a loss of several other crucial PRRs including certain cell membrane-bound TLRs (TLR1, TLR2, TLR4 and TLR5), RIG-I and NOD2 (5, 7, 12–14). This significant gene loss suggests that the intracellular mechanisms in mediating MAMP-PRR activation are likely of extra necessity and importance (5, 13). However, the underlying mechanisms of this process are not fully understood. In this sense, short-read deep sequencing like Illumina RNA-Seq have produced extensive atlases of transcriptomes. In addition, long-read sequencing such as PacBio (yielding reads with average > 15 kb, up to 100 kb or even longer), have proven able to generate high contiguous long reads spanning repeats, and is progressively used in genome assembly and full-length transcriptomics (7, 14, 17, 18). Therefore, integrating both the short-and long-read sequencing approaches would ensure a more precise point-to-point interpretation of the response to immune challenges, especially in organisms with highly complex expanded gene families like Atlantic cod.

Here, we have used Atlantic cod macrophages to further elucidate the functional outcome of the codfish PRR repertoire. Macrophages play a pivotal role in host defence, both phagocytizing non-self-agents and orchestrating subsequent innate and adaptive immune responses (19–21). As a consequence, tissue macrophages perform a critical immune surveillance role with their befitting diversity of PRRs (21). *Ex-vivo* Atlantic cod macrophage cultures were established and challenged with a set of MAMPs mimicking bacterial and viral infections. In addition, a set of well-known pharmacological signalling pathway blockers were used to further unravel cod PRR signalling pathways. Finally, using long-read sequencing technology, a comparison between the predicted transcriptome and the actual macrophage transcriptome was performed. We find that Atlantic cod macrophages efficiently engulf microbes, produce inflammatory mediators and generate a MAMP-specific immune response that agrees with the overall vertebrate models. Sensing of bacterial MDP and of the viral mimic dsRNA is mediated by the NLR and RLR pathways respectively. Comparative analyses demonstrate a role for the internalization for exogenous ligand through scavenger receptors for MAMP-PRR interactions. Intracellular pathways identified through inhibition highlights a central role of TBK1 (TANK Binding Kinase 1) in signal coordination for NOD-based signalling. Thus, our findings demonstrate that the loss of indispensable immunological components, including specific membrane-bound TLRs, does not hamper Atlantic cod macrophage functionality. In contrast, Atlantic cod appears to rely more on cytosolic MAMP sensing using NLR and RLR-based signalling supported by neo-functionalized PRRs.

## Results

### *Ex vivo* cod macrophage culture

Adherent cells derived from cod pronephros and spleen differentiated into a homogeneous macrophage phenotype, with filiform pseudopodia, after three days in culture. Flow cytometry assessment of the cell population demonstrated homogeneity (92%) and was gated for further functional assays (gate R1 in Fig.1A-B, Table S1). Phagocytic capacity was tested with fluorescence-labelled microbes (FITC *Escherichia coli* and *Saccharomyces cerevisiae*) demonstrating 40.77 ± 26.54% (mean ± std. deviation) internalisation of yeast within 1 hour of incubation at 12 °C (Fig.1C and D, Fig S1 and Table S1) and 66.42 ± 3.21% internalisation of *E.coli* after 3 hours of incubation (Fig.1E and F, Fig.S1 and Table S1).

**Fig. 1.**
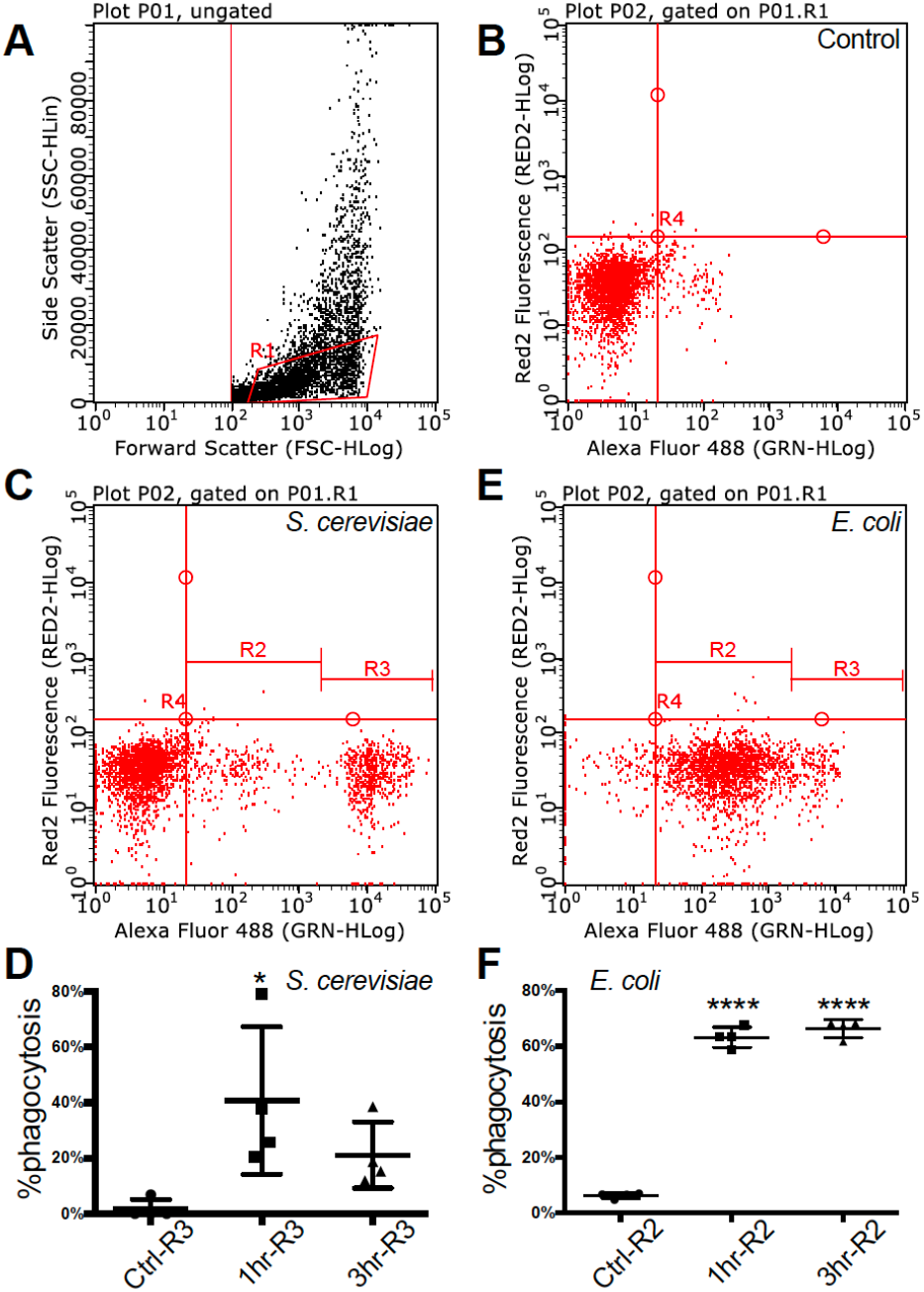
Flow cytometry of fully functional Atlantic cod macrophages during phagocytosis. **(A)** Scatter plot of cell complexity (y-axis) and size (x-axis) that shows the presence of two cell subpopulations: one of morphologically similar cells (R1) and another one of heterogeneous cells with regards to both parameters. **(B, C, D)** The first subpopulation was gated, and it is shown in an untreated resting state **(B)**, during phagocytosis of yeast **(C)**, and during phagocytosis of bacteria **(D)**. **(C’, D’)** Macrophage abundance by size during an untreated resting state, after a 1-hour and after a 3-hour activation of yeast **(C)** and bacteria **(D)**.

### The antiviral response is conserved in the absence of RIG-I

To elucidate the bacterial and viral transcriptional response of the *ex vivo* macrophage culture, a collection of MAMPs were used to mimic a bacterial (‘Bac’, LPS, PGN, CpG DNA) and viral (‘dsRNA’, HMW dsRNA) infection. In total, 101 up-regulated and 4 down-regulated genes were identified for group ‘Bac’ (Fig.2A), and 688 up-regulated and 105 down-regulated genes for group ‘dsRNA’ (Fig.2B). Analysis of annotated DEGs from both challenge groups revealed 46 up-regulated genes expressed in both treatments (Fig.2C, Table S2, Data S1).

**Fig. 2.**
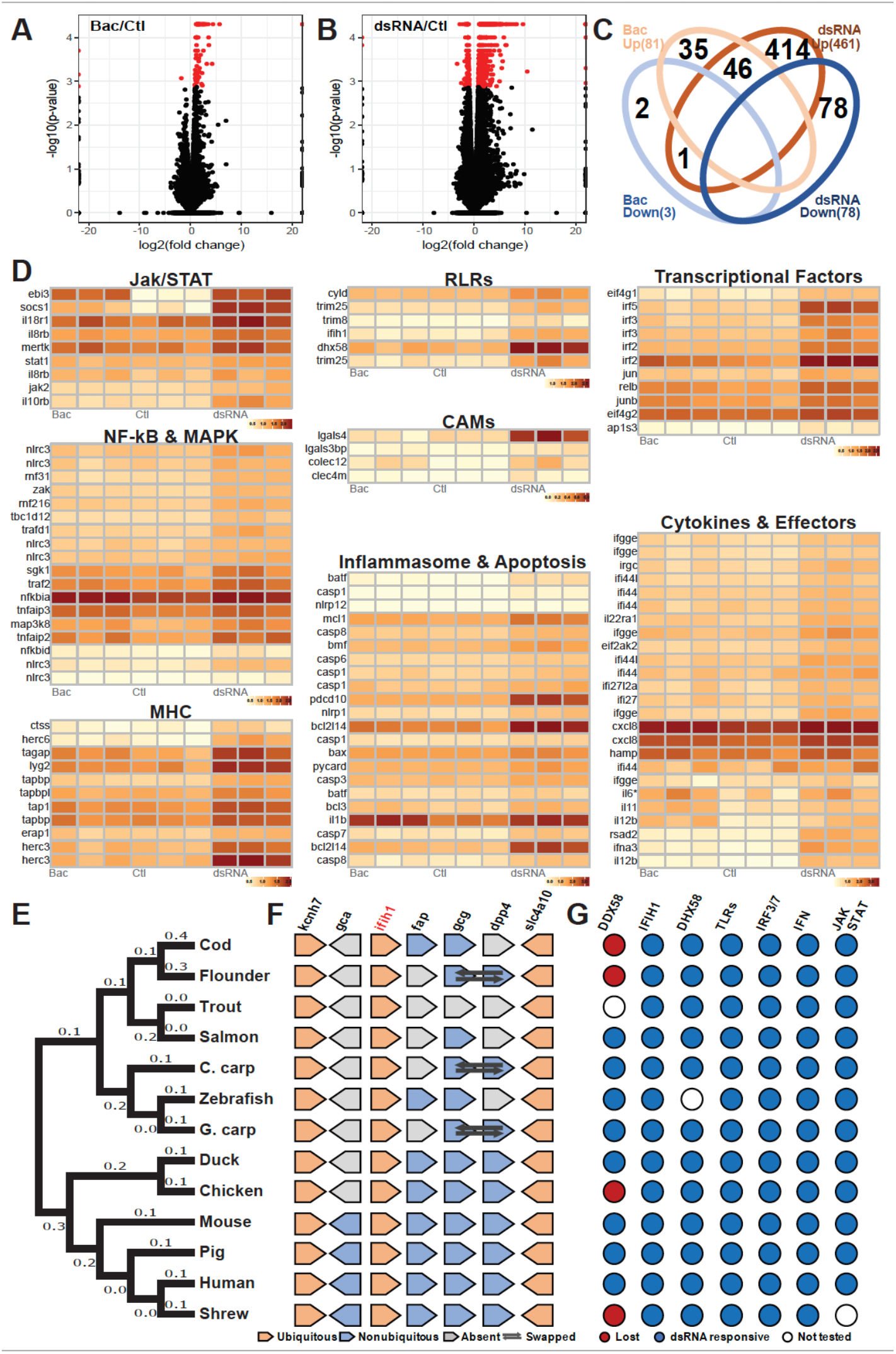
Evolutionarily conserved antibacterial and antiviral response of cod macrophages. **(A, B)** Volcano plots of the expression patterns and statistical significance (shown in red, FDR < 0.05) of genes across three biological samples challenged with **(A)** bacteria-derived MAMPs (Bac) and **(B)** dsRNA contrasted with control samples (Ctl). **(C)** Venn diagram showing number of overlapped up-regulated (UP) and down-regulated (DOWN) genes that had been annotated among Bac and Plc. **(D)** Transcript abundance (log_10_FPKM+1) of selected DEGs, grouped based on shared signalling pathways. **(E)** Phylogenetic tree of *ifih1* from 13 vertebrates. Number of nucleotide substitutions per site are shown next to each branch. **(F)** Synteny analysis of *ifih1*. Pentagon shapes represent each gene, pointing in the direction of transcription. Ubiquitous genes are orange, absent genes are grey, and non-ubiquitous genes are blue. Dark arrows indicate swapped genes. **(G)** Antiviral responsiveness of the “RLRs-TLRs-IRF3/7-type1IFN-JAK/STAT” axis in 13 vertebrates, as shown by published studies. Red circles indicate lost genes, white circles untested genes and blue circles responsive genes.

These were mostly related to membrane internalization and trafficking, activation of transcription and cellular signalling cascades including NF-KB and MAPKs pathways, inflammation-related cytokines, and feedback loop regulators (Fig.2D, Table S2 and Data S1). DEGs (492 total) specific to dsRNA were classified into eight major gene clusters conservatively associated to the antiviral immune response including: i) Lectins; ii) RLRs pathway; iii) transcriptional factors; iv) NF-kB and MAPK signalling; v) JAK/STAT pathway; vi) antigen presentation by MHC class I; vii) inflammasome and apoptosis; viii) Cytokines and effector proteins (Fig.2D, Fig S2, Data S1).

Interestingly, cod, flounder (*Paralichthys olivaceus*)(22), chicken (*Gallus gallus*)(23) and the tree shrew (*Tupaia chinensis*)(24) lack RIG-I (*ddx58*), once considered to be indispensable in jawed vertebrates, and yet can initiate robust antiviral responses. As we observed RLR pathway activation, we looked at other RLR genes in the Atlantic cod genome including *ifih1* and *slc4a10*. Phylogenetic analysis placed the cod RIG-II (*ifih1*) into the teleost lineage as expected (Fig.2E). Further, synteny analysis revealed a conserved genomic gene array structure where *ifih1* is located, with flanking genes *kcnh7* and *slc4a10* being ubiquitously presented across all tested vertebrate genomes (Fig.2F, Table S3). Synteny in the teleost lineage was less conserved for *ifih1*-adjacent genes including *fap*, *gcg* and *dpp4,* which displayed significant variation. Notably, these flanking genes were absent from the trout genome which might be due to the incompleteness of the genome assembly and/or genomic rearrangement. Moreover, a survey of functional studies in these 13 vertebrates using dsRNA (or in some cases viruses) as stimuli highlights a conserved antiviral signalling axis, where key components RLRs, TLR3, IRF3/7, type I IFN and JAK/STAT are involved (Fig.2G, Table S4).

### Muramyl dipeptide drives bacterial MAMP recognition

In fish, the recognition of bacterial MAMPs and their respective roles in downstream signalling and activation remains under debate (25). In cod, the absence of homologs of the major cell surface mammalian TLRs allowed for further exploration of the downstream signalling pathway following LPS, PGN and CpG DNA detection. Using *il1b* and *il6* mRNA expression as markers of MAMP recognition, a subsequent pro-inflammatory response, although non-significant, was shown in macrophages to LPS (Fig.3A) and CpG DNA (Table S5). LPS signalling across the Teleostei remains unclear with the majority of fishes lacking TLR4 however the recent reporting of *caspy* as an intracellular LPS receptor in zebrafish suggests an alternative route for LPS sensing(26). Ultra-pure PGN (MDP; muramyl dipeptide) treatment induced a significant activation both 3- and 12-hours post challenge with MAMPs, while peptidoglycan did not (Fig.3A and Table S5)

**Fig. 3.**
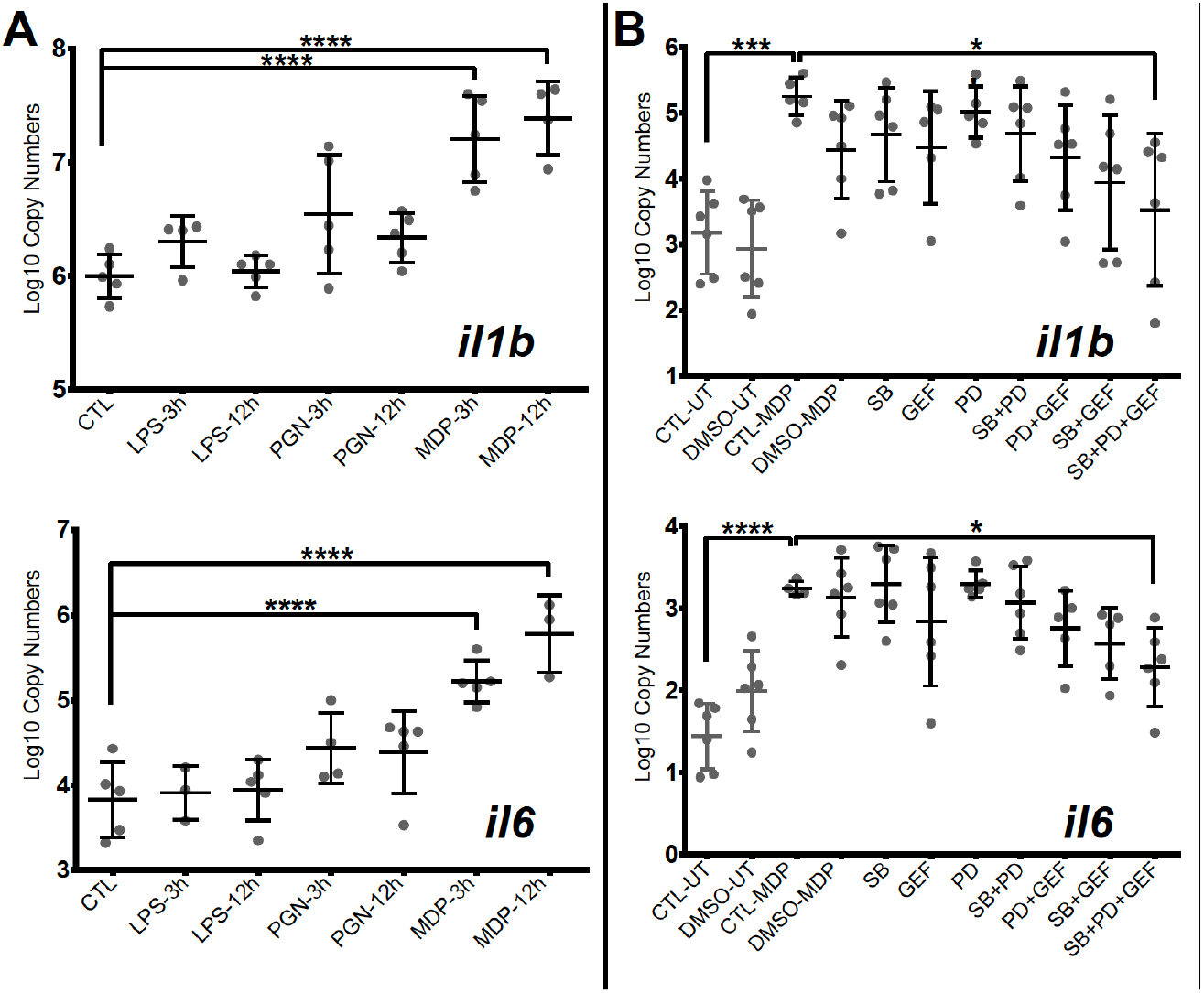
Muramyl dipeptide peptidoglycan (MDP) triggers the strongest antibacterial response in Atlantic cod macrophages, mediated by NOD1 and MAPKs. **(A, B)** Copy number (log10) of interleukin *il6* and *il1b* in response to **(A)** an activation by various individual MAMPS during 3 and 12 hours and **(B)** inhibition of NOD pathway-relevant genes in MDP-activated samples. PD: MEK1 and MEK2 Inhibitor, GEF: gefitinib, SB: p38 MAP Kinase Inhibitor. Whiskers show the standard deviation. Asterisks represent statistical significance: *<0.05, ***<0.001,**** < 0.0001. Abbreviations: CTL: control, LPS: lipopolysaccharide, PGN: peptidoglycan, MDP: Muramyl dipeptide peptidoglycan, UT: untreated, DMSO: dimethyl sulfoxide

To further delineate the signalling pathways involved, we employed a suite of kinase inhibitors targeting p38 MAPK and MAPKK (MEK1 and MEK2), including a NOD1/2 inhibitor acting through RIPK2. None of the inhibitors significantly blocked MDP-driven *il-1b* and *il-6* mRNA increases while used individually. However, when combined, a significant inhibition was observed (32.9% for *il-1b* and 29.5% for *il-6*) (Fig.3B and Table S5). Although sequence divergence may impact upon cod-specific MAPK inhibition, combinations of RIPK2 and MAPK inhibition produced a more pronounced effect in comparison to MAPKK and p38 inhibition.

MDP-treated macrophage cultures that were positive for both *il1b* gene expression (Fig.4A) and extracellular secretion of the pro-inflammatory mediator prostaglandin E2 (PGE_2_) (Fig.4B) were analysed by RNA-Seq. MDP-activated macrophages showed higher activation (414 DEGs) than in response to the previously used combination of bacterial MAMPs (LPS, PGN and CpG DNA) (105 DEGs). Interestingly, at 3 hours post MDP treatment only 5 up-regulated DEGs and no down-regulated DEGs were observed (Fig.4C), while 286 up-regulated and 123 down-regulated DEGs were identified after 12 hours of treatment (Fig.4D, Fig S3 and Data S1). The majority of DEGs (90%) were annotated, and functionally categorized based on six major pathways: i) G-protein-coupled receptors (GPCRs) and endocytosis; ii) PRRs sensing; iii) apoptosis; iv) NF-kB and MAPK; v) cytokines; and vi) transcription factors (Fig.4E, Data S1). Both *il1b* and *ripk2* were significantly regulated consistently with previous gene expression results.

**Fig. 4.**
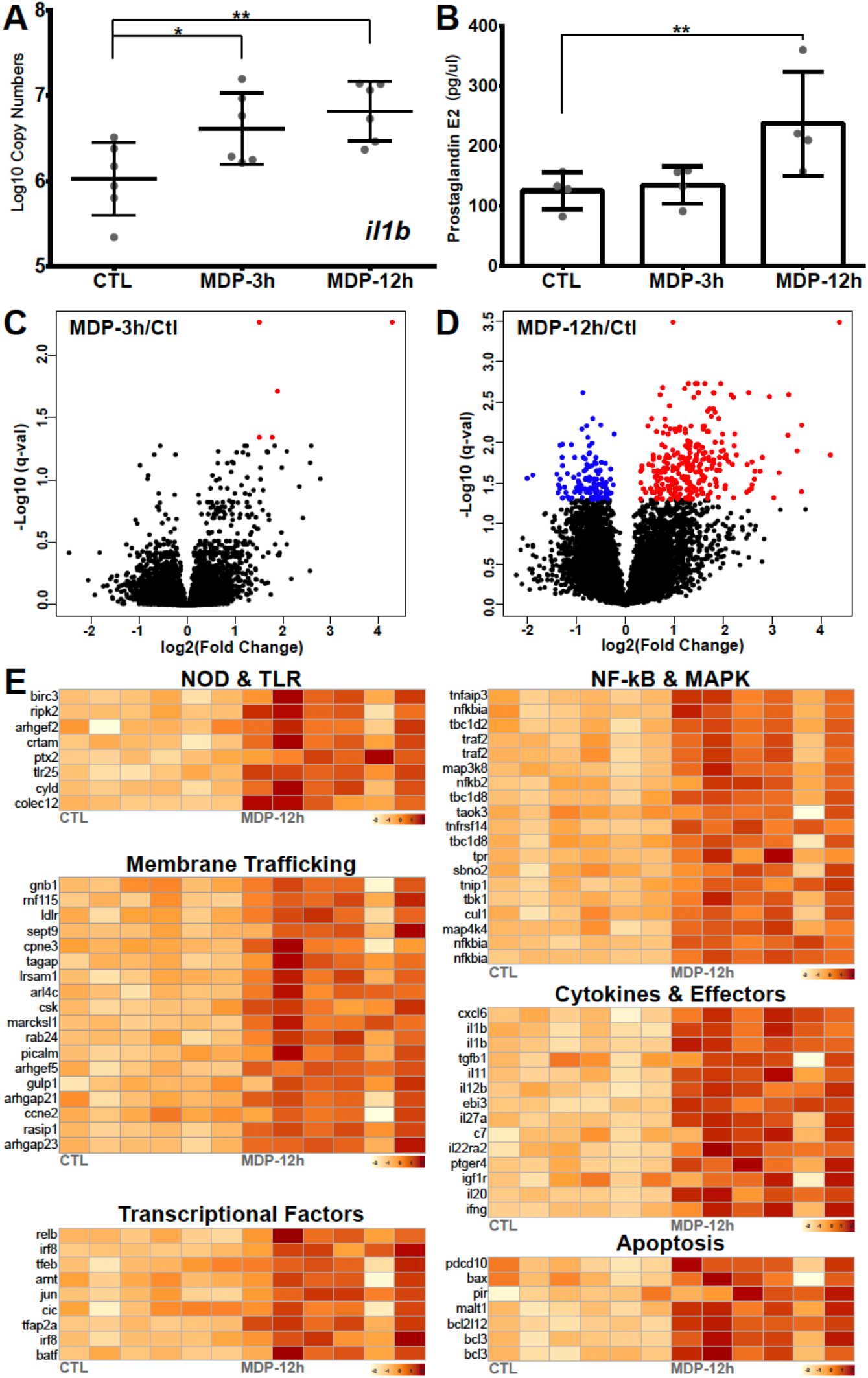
Antibacterial response to MDP challenge. **(A, B)** Immune response to Muramyl Dipeptide (MDP) measured by **(A)** copy number (log10) of *il1b* and **(B)** prostaglandin E_2_ secretion in *ex-vivo* cultured cod macrophages. **(C, D)** Volcano plots of the expression patterns and statistical significance (red if significantly up-regulated, blue if significantly down-regulated, FDR < 0.05) of genes across six biological samples challenged with MDP for **(C)** 3 hours and **(D)** 12 hours contrasted with control samples. **(E)** Transcript abundance (log10FPKM+1) of selected DEGs challenged with MDP for 12 hours. Grouped based on shared signalling pathways.

### Divergent MAMP-PRR recognition systems converge

The extensive repertoire of MAMP sensing in animals is underpinned by a diverse set of adaptors and signalling proteins that form pathways that ultimately activate the inflammatory response. In cod, dependence upon intracellular MAMP sensors may be expected to impact upon the activation of critical signalling pathways. Using GO enrichment analysis with cod DEGs annotated to human SwissProt identifiers, significantly enriched KEGG pathways for dsRNA- and MDP up-regulated DEGs were shown to be partially shared (Fig. 5 and Data S1). Six major functional clusters with their associated DEGs highlighted extensive convergent signalling in cod macrophages upon distinct MAMP challenge (Fig. 5 and Fig. S5), consistent with shared DEGs between dsRNA and MDP (Table S2).

**Fig. 5.**
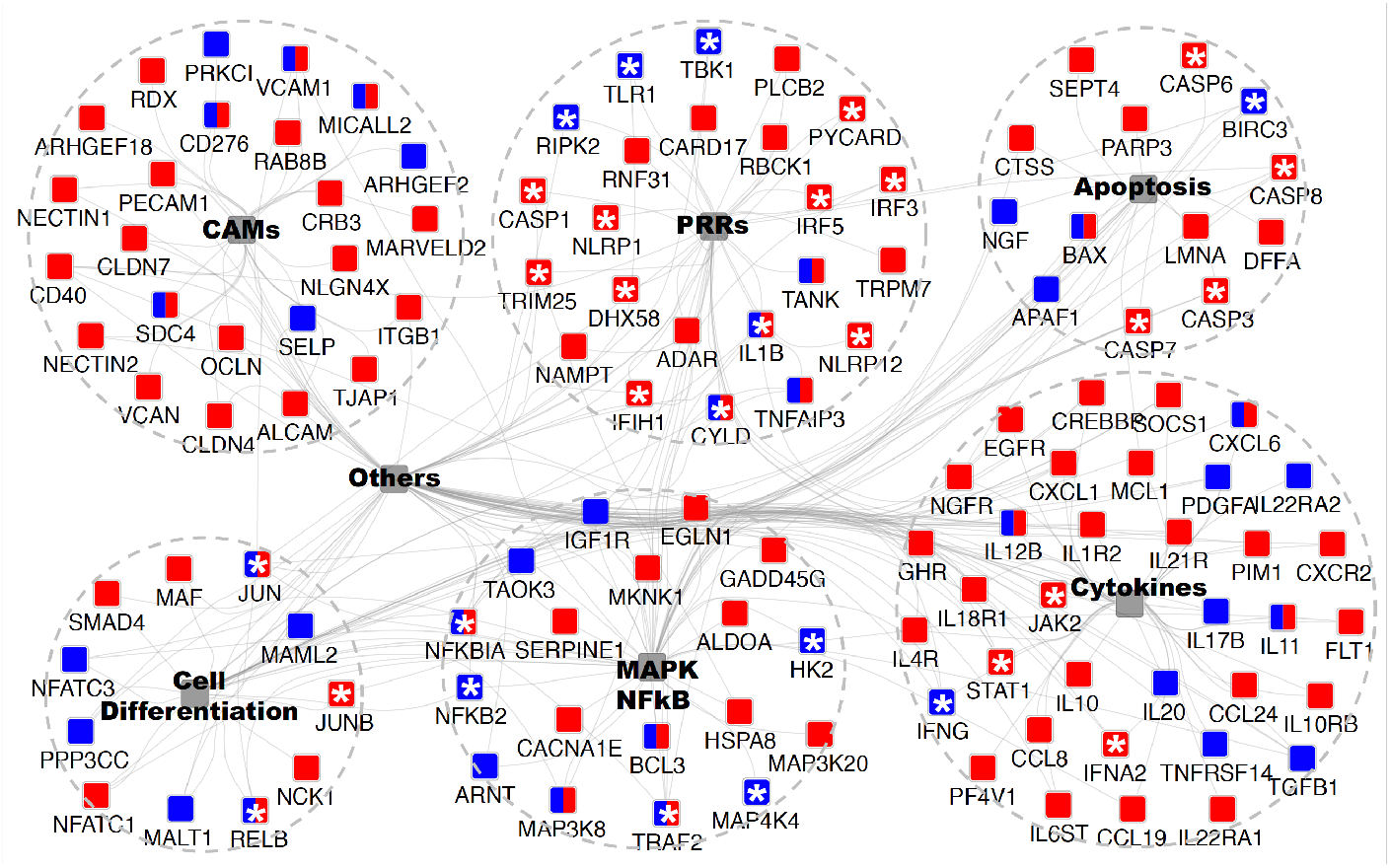
GO enrichment analysis of MAMP up-regulated DEGs to identify underpinned functional pathways in activating the cod macrophage inflammatory response. GO enrichment and network visualization was conducted by Cytoscape plugin ClueGO and CluePedia. Cod up-regulated DEGs annotated to human SwissProt identifiers were mapped to KEGG (Kyoto Encyclopedia of Genes and Genomes) pathways database. Enriched KEGG terms that are shared the same cluster were collapsed into one integrative node (shaded grey) labelled with brief descriptions respectively (See Fig. S5 and Dataset.S1 for the full network). Six major inflammation-related functional pathways were shown, and the remaining ones were labelled with “Others”. The associated DEGs with indicated KEGG pathways were colour coded in a MAMP-dependent fashion, red and blue nodes denote dsRNA- and MDP-responsive genes, respectively; whilst the dual-labelled nodes denote the common DEGs that are responsive to both MAMPs. Asterisks indicate genes that serve as key sensors, adaptors or regulators in mediating inflammatory response and had been intensively discussed in texts.

It is also noteworthy that the MAMP sensing pathways, for both dsRNA and MDP, in cod demonstrate stimuli-specific DEGs that converge upon common thematic regulators, thus providing diversity in MAMP-specific regulators of signalling. For dsRNA-specific pathways, RLR- and NLR-receptors, JAK-STAT signalling, IRF3/7 signalling, type-I IFN response and MHC class I antigen presentation pathways were shown to be conserved across vertebrates, from mammals (27) to teleosts (28, 29) (Data S1 and Fig. S5). In contrast, the cod MDP-specific pathway demonstrates a different MAMP sensing mechanism where NOD signalling, pro-inflammatory responses, type-II IFN responses and leukocyte differentiation mediated by *IFNG*, *IRF8* and *TGFB1,* are enriched.

### Evidence for (sub) neofunctionalisation in cod PRR

To further capture the full-length mRNA isoforms, especially those of significance in MAMP sensing, long read PacBio transcriptome profiling (IsoSeq) was performed. After polishing by Quiver, 34,516 high-quality (hq) and 157,657 low-quality (lq) transcripts were obtained. Merging of the two, resulted in 192,173 isoforms with average length of 2,351 bp were obtained. In total, 69.75% of the transcripts were uniquely mapped to the cod genome assembly (gadMor2). For some of the low expressed genes detected using RNA-Seq (e.g. FPKM< 1), such as *colec12*, *nlrc3* and *ifih1* (RIG-II), long reads were not detected, probably due to low sequencing coverage. Most cod PRRs found in the Illumina sequencing data were captured at least once by IsoSeq. Inspection of the full-length transcripts of these PRRs obtained by IsoSeq validated the high sensitivity of short reads assembler Cufflinks and StringTie (Fig.S5). For instance, three potential transcripts for RIG-III (*dhx58*) were assembled by Cufflinks, of which only one isoform “TCONS_00018715” was captured by IsoSeq (Fig.S6). Accordingly, the most dominant (>75%) isoform “TCONS_00018715” had the highest abundance (FPKM=21.06) in untreated resting macrophages (Ctl). Additionally, two other isoforms “TCONS_00018716” (with an extra 5’-end exon), and “TCONS_00018717” (with exon skipping) were dsRNA-challenge specific. Multiple transcripts for *tlr25* were identified by StringTie (after filtering out transcripts with low variance (see Supplementary Information for details), of which only transcript “MSTRG.16445.4” were significantly activated by MDP (Fold Change=2.831, FDR=0.016). Sequence alignment of all these transcripts towards *tlr25* isoforms (a~g) from Solbakken et al.(12, 13), showed that the isoform “MSTRG.16445.4” is identical to *TLR25d* (13)(Fig.S6).

## Discussion

Experimental studies, prior to the discovery of the alternative arrangement of the Atlantic cod immune system (5, 30, 31), described the Atlantic cod immune response, including survival (32), fitness (33), pathogen elimination (34), and antibody-production (35), as being broadly similar to other teleost fish. Furthermore, the loss of most membrane-bound TLRs, MHC II, CD74 invariant chain and the CD4 receptor does not appear to have imposed restrictions on the Gadiformes, a highly successful family (>400 reported species)(2). Therefore, a more prominent role, due to increased gene dosage, for intracellular PRRs and MHC I has been proposed (5, 13).

In order to assess the MAMP-PRR response in cod, an *ex-vivo* macrophage cell culture model was developed. As one of the major professional antigen-presenting cells (APCs), highly phagocytic macrophages are responsible for innate-adaptive immunity cross-talk(21). Differentiated cod macrophages efficiently engulf microbes, produce inflammatory mediators (PGE_2_) and generate a canonical time-dependent MAMP-specific immune response in agreement with vertebrate macrophage models (19, 20, 27) (multidimensional scaling; Fig.S2 and S3, functional clustering; Fig.5). The MAMP-activated cod macrophage transcriptome is characterised by profound changes in the expression of cytokines, chemokines, antimicrobial peptides and transcriptional factors (Data S1), driven by a set of stimuli-specific gene-clusters tailored to cope with different MAMP-PRR signals. Consistently, functional interactomes for dsRNA and MDP responsive DEGs are all convergent into functional pathways such as NF-kB and IRFs (Fig.2 and Fig.4). Similar results were reported in *in-vivo* studies on cod using the facultative intracellular bacterium *Francisella noatunensis*(36). Our data demonstrates that in cod, macrophage immune response specificity is generated in a ligand-dependent manner in the absence of certain membrane-bound TLR interactions (Fig.2 and Fig.4).

The co-evolution of both diversity and restriction across the different components of the immune system is an intriguing puzzle, but hampered by the challenge of developing suitable experimental models(37). Strong evolutionary constraints imposed upon metazoan regulatory molecular circuits ensure robust regulatory mechanisms whereas pressure to diversify recognition structures toward pathogens is critical(9). Interestingly, the deletions observed in the cod genome appear contradictory in that specific cell-surface TLR are absent thus reducing recognition capacity(5), although functionality in cod macrophages is not impaired. Studies in Atlantic cod suggest that endocytosis of lipoprotein and polyribonucleotides (dsRNA) is mediated by scavenger receptors(38). For exogenous dsRNA sensing, variations of membrane-trafficking delivery can trigger distinct immune responses(39, 40). Lipid-based dsRNA transfection induces a highly efficient and scavenger receptor Class-A (SR-As) independent endocytosis triggering TLR3-mediated apoptosis(39, 40). In contrast, directly adding dsRNA to the culture medium, as in this study, induces a SR-As-dependent endocytosis, mediated by the RLR family of cytosolic receptors(39, 40). In this study, cod macrophages increased the transcription of the Scavenger Receptor Class-A Member (SR-As) *colec12* (aka SCARA4) with both dsRNA and MDP indicating activation of SR-As-dependent endocytosis. Moreover, many other vesicle-trafficking related pathways were significantly activated indicating a common response (Fig.5 and sData 1). In accordance, elevated transcription of all cod RLRs, including *ifih1* and *dhx58*, post dsRNA challenge was observed. In contrast, no activation was observed for *tlr3* or its relevant adaptors including *myd88* and *trif*. PGN does not induce any inflammatory response via direct microinjection into cytosol in mammals, unless delivered by membrane encapsulated-vesicles(41). In fact, recent studies highlight the endosome as a crucial platform for NOD1/2-dependent ligand sensing and inflammatory signalling(42–44). Despite *nod1* not being significantly induced by MDP (Fold-change: 1.7; *p*-val: 0.01; FDR: 0.13, Fig.S4), transcription of its adaptor (*arhgef2*), enzymes (*ripk2, hk2*), and ubiquitination regulators (*birc3*, *cyld*, *tnfiap3*) were all consistently elevated in our study which is consistent with observations from studies in mammals(43, 45). Additionally, results from our blockade assays using a cocktail of inhibitors against RIPK2 and MAPKs further validated the role of NOD-pathways in sensing MDP in cod macrophages. Therefore, despite the absence of *nod2*, the NOD pathway in cod remains functional, in line with observations of NOD1-mediated antibacterial responses in birds lacking NOD2 including both duck(46) and chicken(47).

In addition to the loss of specific TLRs and the classical MHC II pathway, some intracellular PRRs including RIG-I and NOD2 were also found to be absent in the Atlantic cod(13). Similarly, RIG-I is lost in the coelacanth, spotted gar, cave fish, lamprey and tree shrew and NOD2 is absent in the spotted gar, frog, reptiles and birds (Ensembl Gene gain/loss tree). We found that the remaining MDA5/LGP2 and NOD1 initiate the RLR- and NOD-pathways specific to dsRNA and MDP sensing, respectively, in Atlantic cod. In the tree shrew, MDA5 and LGP2 synergistically mediate dsRNA sensing in compensation for the loss of RIG-I(24). This is achieved by the RIG-I-specific adaptor, STING (Stimulator of interferon genes protein, encoded by *tmem173*), that acquired a novel MDA5 interacting function(24). In zebrafish, NOD1 knockout mediated by gene-editing dramatically affected many other NLRs members by inducing expression whilst conventional NF-κB and MAPK immune signalling pathways were unimpaired(48). Interestingly, in line with above mentioned PRRs, in dsRNA or MDP challenged cod macrophages, other receptors such as *nlrc3s*, *nlrp1*, *nlrp12* and *tlr25* were also significantly up-regulated (Fig.5 and SData 1). In contrast to mammals, the dramatically expanded NLR and TLR gene repertoire in the cod genome suggest a potential rapid functional shift(7, 14). TLR25, one of the teleost-specific TLRs, belongs to the TLR1 family and shares significant sequence similarity with mammalian TLR1(12, 13). In this study, one of the TLR25 isoforms, *tlr25d*, was uniquely regulated by MDP challenge, however no changes were observed in any Myddosome or Triffosome related factors leaving its downstream signalling in cod unclear. On the other hand, several NLR family members’ associated adaptors including *pycard* and *casp1* were induced suggesting that these mediators of inflammasome formation are active in Atlantic cod as reported in mammals(49). Recent studies, in zebrafish, described a conserved functional role for the NLRP1 inflammasome in teleosts(50) and our observations further support a central role for NLRs in innate immunity in the Atlantic cod and across the Teleost fish.

The underlying molecular patterns of activation in cod macrophages point toward a conserved cytosolic MAMP-PRR recognition that link to supra molecular organising centres which themselves appear to be conserved across vertebrate innate immune signalling(9, 29). RLRs, NLRs and TLRs engage in convergent signalling cascades to regulate the expression and degradation of pro-inflammatory mediators upon ligand sensing, that are highly conserved owing to strong functional and regulatory constraints(51, 52). Interestingly, in fish canonical LPS-TLR4 sensing is seemingly absent(53), with the exception of zebrafish, as a negative regulator of inflammation(25). However, recently caspy2 has been reported as an intracellular LPS receptor in the zebrafish(26). In this study, no significant response to a standard LPS preparation was detected. However, a robust MDP-NOD driven activation of inflammation coupled to an enrichment of the IRF8 pathway was observed (Fig. S5 and Dataset S1). In mice, IRF8-impairment led to reduced secretion of IL-12, an essential inducer of T helper (Th1) cell polarization, causing a dramatic increase in susceptibility to intracellular infections highlighting a critical defence function(54). Furthermore, a regulatory role for IRF8 has been suggested in MHC II antigen presentation(55, 56). Recently, MHC class I based cross-presentation through the acquisition of functional sorting motifs has been suggested as a mechanism in cod for MHC II loss(57). The cod macrophage model and further exploration of the IRF8 pathway provides a platform for further functional studies. Multiple genome-based studies across a wide range of fishes highlight an exceptional diversity at the level of cytosolic PRR families(58–60). It is tempting to speculate that the environment, both aquatic and internal, coupled to restricted adaptive immunity has driven innovation in cytosolic MAMP detection.

Available evidence, including this study, suggests that functionality including intracellular PRR systems are conserved throughout vertebrates, and thus, also provide an efficient host defence in the Gadiformes lineage contributing to their evolutionary success(29, 36, 61, 62). In light of the loss of the majority of extracellular PRR capability and lack of MHC class II-based antigen presentation it suggests, at least in the codfishes, these functions may not be ‘core’ to the immune response and possibly bony fishes in general. Notably, evidence for sub- or neofunctionalisation of PRR, in a ligand-dependent manner, is demonstrated via several NLRs paralogues and one of the TLR25 isoforms. Sub- or neofunctionalization following gene duplication of multigene gene families driven by selection is a recognized mechanism of genome innovation and adaptation(4). The dramatically diversified NLR and TLR paralogues, coupled with their complex protein-protein interactions via ligand-sensing and signal transduction domains, presumably allow the cod immune system to adapt to fast-evolving pathogens.

In summary, our results, based upon a novel *ex vivo* macrophage cell culture, demonstrate that intracellular PRR systems based upon RLR and NOD signalling, in the absence of RIG-I and NOD2, are critical drivers of the response to bacterial and viral pathogens in Atlantic cod. Our analysis highlighted the inflammatory response to be intimately linked to scavenger receptor-based internalization of MAMPs coupled to highly conserved intracellular signalling platforms. Cytosolic PRR capabilities in the Atlantic cod are functional, in line with most vertebrate studies, and are based upon canonical NF-kB and IRFs activation pathways. Additionally, we uncover that the multiple NLR paralogues and the unique TLR25 isoform identified in Atlantic cod display ligand-dependent differential expression suggesting (sub)neofunctionalisation toward specific immune defensive strategies. Our results further demonstrate that the extreme remodelling of the Atlantic cod immune system provides an unprecedented opportunity to explore the evolutionary history of PRR-based signalling in vertebrate immunity.

## Material and methods

### 1. Samples

One-year old Atlantic cod, *Gadus morhua* L., reared in land-based tanks supplied with filtered sea water were obtained from Ardtoe Marine Facility (UK) during 2014-2015. They were kept at environmental temperature and fed on commercial dry pellets once a day. All procedures were in accordance with UK Home Office welfare guidelines. The fish were sacrificed with UK Home Office approved methods, using an overdose of the anaesthetic benzocaine (Sigma-Aldrich) followed by brain concussion. The head kidney and spleen of each animal were dissected out and kept at 4 °C in high glucose Dulbecco’s Modified Eagle Medium (DMEM) (Sigma-Aldrich) and 0.2% Primocin™ (InvivoGen) until cell culture.

### 2. Macrophage cell culture

The sampled tissues were ground together through a 100 μm cell strainer (Fisher Scientific) in a proportion of 1:3 spleen to head kidney into a tube with fresh DMEM solution with 0.2% Primocin. Cells were precipitated by centrifugation at 1500 rpm for 5 minutes and resuspended in DMEM (0.2% Primocin and 10% chicken serum (Life Technologies)). The cell culture plates were coated with 0.5 ml of poly-D-lysine (Sigma-Aldrich) (0.1 mg/ml in no calcium, no magnesium Dulbecco's Phosphate-Buffered Saline (DPBS) (Life Technologies)) per well for 30 minutes, rinsed with 1.5 ml of DPBS per well, and left to be air-dried prior seeding. Twelve-welled plates (Thermo Scientific) were filled with 1 ml per well of the resuspended cells (which contained approximately 2 million cells per ml). The cell cultures were kept in an incubator at 15°C with 4% CO_2_ supply. After one day and thereafter every two days (5 days in total), half of the medium per well was replaced with fresh DMEM (0.2% Primocin and 10% chicken serum from the same batch).

### 3. Phagocytosis assay

Alexa fluor 488 labelled bacteria (*E. coli*, Sigma-Aldrich) and yeast (*S. cerevisiae*, Sigma-Aldrich) were used as phagocytosis targets. After 1 hour and 3 hours of incubation with bacteria and yeast respectively, macrophages from four individuals were analysed using Guava^®^ easyCyte™ 8HT Flow Cytometer (Merck Millipore) to measure phagocytosis. Flow cytometry assessment of the cell population demonstrated homogeneity (92%) and was gated for further functional assays (gate R1 in Fig. 1 *A-B*).

### 4. MAMP Immune challenge

One hour prior to stimulation, the cell culture medium was replaced with serum-free DMEM, eliminating the possible interference of serum with the assay. Ten μl of 1mg/ml stock solutions of MAMPs (i.e. Lipopolysaccharide (LPS; LPS-EK*;E.coli*), Peptidoglycan (PGN; PGN-EK*;E.coli*), Muramyl Dipeptide (MDP; PGN-*E.coli* ndss ultrapure), dsRNA (High Molecular Weight, HMW) and CpG (*E. coli* ssDNA/LyoVec)) per ml was used in all cases to challenge the cells for 3h and 12h (Table S6). All MAMPs were purchased from Invivogen.

### 5. Inhibition of the immune response to MDP

MAPK Kinase Inhibitors PD (MEK1 and MEK2 Inhibitor, Sigma-Aldrich), SB (p38/ERK MAP Kinase Inhibitor, Sigma-Aldrich), and NOD1/2 inhibitor GEF (Gefitinib, Invivogen) were added to the cell cultures along with either solution buffer (control) or MDP (10 μg/ml). The working concentrations of the inhibitors were, respectively, 2, 0.5 and 10 μM (Table S6). The cultures were incubated for 12 hours. All possible combinations of inhibitors were tested.

### 6. Prostaglandin E_2_ measurement

Supernatants (serum-free DMEM with 0.2% primocin) of control and MDP-challenged macrophages from 4 different individuals were preserved at −20 °C prior testing. Prostaglandin E_2_ (PGE_2_) was quantified by a monoclonal EIA kit following the manufacturer instructions (Cayman, USA).

### 7. RNA isolation and quality control

After the whole supernatant had been removed by pipetting, 250 μl of TriReagent^®^ (Sigma-Aldrich, UK) was added to each well of a 12-welled plate. Total RNA of the macrophages was extracted by the phenol/1-bromo-3-chloropropane method (Sigma-Aldrich, UK) following the manufacturer’s instructions, with 1 μl of glycogen (Roche) per 1 ml TriReagent^®^ added to enhance the RNA precipitation during the isopropanol step. The total RNA samples were then subjected to NanoDrop (Thermo Scientific) to determine the RNA concentration, and Bioanalyzer (Agilent Technologies) to estimate the integrity and purity of the RNA samples. Only the samples with RIN values greater than 8 were kept. Qubit^®^ (Life Technology) was used to quantify the concentrations of the samples that were going to be sent for sequencing. All samples were stored at −80°C.

### 8. cDNA synthesis

Different cDNA synthesis protocols were used depending on the purpose of the samples, following the manufacturer’s instructions of each kit. The Illumina RNA-seq cDNA library was prepared using 1 μg of RNA from each sample and TruSeq V2 kits (Illumina, CA, USA), with reduced RNA fragmentation time (3 minutes) to maximize obtention of longer reads. For PacBio IsoSeq, 1 μg of RNA per sample was used for first-strand synthesis by SMARTer PCR cDNA Synthesis Kit (Clontech). For quantitative RT-PCR (qPCR), 1 μg of total RNA per sample was used to synthesize cDNA with SuperScript III RNase Transcriptase (Invitrogen) and Oligo-dT primer (Invitrogen).

### 9. Library preparation and sequencing

Two rounds of Illumina RNA-seq were performed. For the first one, which consisted of a preliminary MAMP activation screening, RNA samples from macrophage cultures derived from 3 fish were pooled. In total, 9 pooled libraries were prepared: 3 control pooled samples (Ctl), 3 bacterial MAMP-activated pooled samples (Bac) and 3 dsRNA-activated pooled samples (Plc) (Table S7). They were loaded onto one lane of an eight-laned FlowCell Chip. The second round looked at the effects of exclusively MDP on the cod macrophage transcriptome. For these, a total of 18 libraries were prepared, each individual fish had three aliquots of one control sample, one sample challenged with MDP for 3 hours and another one challenged with MDP for 12 hours. These were loaded onto two lanes of an eight-laned FlowCell Chip.

To prepare the libraries for IsoSeq, the Pacific Biosciences IsoSeq library preparation protocol was followed. Before library preparation, the cDNA was first size-selected using BluePippin (Sage Science) and then using AMPure^®^ XP (Auto Q Biosciences) beads. Then they were sequenced on a Pacific Biosciences RS II instrument using P6v2/C4 chemistry (Pacific Biosciences of California Inc., Menlo Park, CA, USA). In total, 2 SMRT cells for 1-2kb, 3 SMRT cells for 2-3kb and 4 SMRT cells for 3-6kb were used for sequencing.

### 10. Absolute quantitative RT-PCR

All primers for qPCR were designed using BatchPrimer3 v1.0 based on the target genes annotated in the latest cod genome assembly gadMor2 (Table S8) (14). Target genes were validated using thermal gradient RT-PCR and the products that met the quality criteria were cloned into bacterial plasmids. Two micrograms of cDNA were used as a template for PCR with gene-specific primers. Target mRNAs were amplified using MyTaq HS DNA Polymerases (Bioline, UK), and amplicons were run on 1% agarose gels, stained with ethidium bromide and purified with NucleoSpin^®^ Gel and PCR Clean-up (Macherey-Nagel, Germany). Purified PCR products were ligated in pGEM-T easy vectors (Promega, USA) and transformed into *Escherichia coli* (DH5a strain). One selected transformant of each construct was grown to obtain plasmid DNA (Miniprep kit, Macherey-Nagel). All constructs were verified by Sanger DNA sequencing with T7 and SP6 primer sets (GATC Biotech, Table S8). Pro-inflammatory cytokines mRNAs for *il1b* and *il6* were selected on account of their high inducibility during PAMP responses (Fig.S7). See details for PCR programme and reaction systems in supplementary Information. One-way ANOVA was used to test the statistical differences between indicated experimental contrast. Significance was reported if *P* < 0.05. Graphs were plotted using Prism6^®^ (Table S5).

### 11. Transcriptome assembly and transcript abundance estimation

Raw Illumina reads from each sample were trimmed using Cutadapt (v1.4.2) with cutoff as Phred < 20. For the downstream differential gene and transcript expression analysis two pipelines were used: (1) reference genome based using the Tuxedo pipeline (Tophat2, cufflinks and cummeRbund) (63): Trimmed reads from macrophage samples (3x Ctl, 3x Bac, 3x Plc) were mapped to the gadMor2 cod genome assembly (14) with the short read aligner Tophat2 (v2.0.9) (see Supporting Information for details). The reads from all samples were assembled by Cufflinks (v2.2.1) and the results were visualized and plotted by R package ‘CummeRbund’. (2) The updated Tuxedo pipeline consisting of Hisat2-Stringtie-Ballgown (64). The new tuxedo pipeline was used to analyse the MDP activated samples,). The trimmed reads from MDP activation with different time-intervals were mapped to the same genome assembly by Hisat2 (v2.0.5), assembled, merged and their expression abundance was estimated by Stringtie (v1.3.2-Pre) with default settings (64). The results were visualized and plotted with the R package ‘Ballgown’.

### 12. Transcriptomic sequence annotation

The nucleotide sequences of differentially expressed genes (DEGs) from the contrast between control and the various treatments were abstracted by Gffread from merged assemblies using Cufflinks and Stringtie (Table S9). The sequences were then annotated based on its similarities with human protein sequences and aligned to the human UniProt database (downloaded in Jan 2017) by Blastx (Blast+ v2.2.29). The non-redundant DEGs were obtained by collapsing the assembled “gene identifier” from both gene and transcript clusters and the unique ones were kept. Subsequently, the overlapping analysis of those DEGs were conducted by Venn Diagram with its annotated human gene names. **Dammit** (v2.0.1) was used to predict open reading frames and annotate from polished IsoSeq transcripts.

### 13. Structure of MDP-responsive PRRs by full-length transcriptome sequencing

All the reads from three size-selected cDNA fractions were clustered and polished by IsoSeq_SA3nUP. After polishing by Quiver, 34,516 high-quality (hq) and 157,657 low-quality (lq) transcripts were obtained. All full-length transcript isoform sequences (with either high- or low-qualities) were merged, yielding 192,173 isoforms with average length of 2,351 bp. In total, 69.75% of the transcripts were uniquely mapped to the cod genome assembly (gadMor2) by aligner Star (2.5.3a) under instructions of IsoSeq_SA3nUP. The alignments of transcripts were sorted by Samtools (v0.1.19) and visualized with the R package ‘Gviz’.

### 14. Synteny of *IFIH1* (RIG-II)

Full-length protein sequences of RIG-II for non-cod species were obtained from NCBI and used directly for phylogenetic analysis. Sequences were first aligned by ClustalW and a maximum likelihood tree was produced using Maximum Likelihood (ML) with 2000 bootstraps by MEGA7. The evolutionary history was inferred by using the Maximum Likelihood method based on the JTT matrix-based model. Synteny analyses of IFIH1 among selected vertebrates are based on genome assemblies from NCBI (RefSeq Release 82), except cod, trout and grass carp whose genome resources are available elsewhere (Table S4).

### 15. Functional Gene Ontology enrichment analysis

Human gene identifiers were retrieved from UniProt-SwissProt for all DEGs. Gene clusters were further divided by different immune stimuli (dsRNA/MDP) and expression patterns (up/down). Each of them was separately provided as input gene cluster and analyzed together. Cytoscape plugin ClueGO and CluePedia were to perform the GO enrichment analysis against KEGG (Fig.5) and comprehensive functional databases including KEGG, GO, WikiPathways and REACTOME_Pathways (Fig. S5 and Dataset S1). The following ClueGO parameters were selected: Go Term Fusion; display pathways that with adjusted-p-values less than 0.05; GO tree interval levels from 3 to 8; GO term minimum numbers of genes (#) and percentage of genes (%): #4+6%, #2+3%, #3+4%, #2+3% for dsRNA-Up, -Down, MDP-Up, -Down respectively; kappa score 0.3; GO enrichment/depletion by two-sided hypergeometric test and corrected by Benjamini-Hocheberg; GO terms are presented as nodes and grouped together based on functional similarity. Node size is negatively proportional to the adjusted *p*-value for GO enrichment; interactome layout by Cytoscape plugin AllegroLayout with “Allegro Spring-Electric” option.

## Acknowledgements

The sequencing service was provided by the Norwegian Sequencing Centre (www.sequencing.uio.no), a national technology platform hosted by the University of Oslo and supported by the "Functional Genomics" and "Infrastructure" programs of the Research Council of Norway and the Southeastern Regional Health Authorities. All bioinformatics analysis was performed using Abel high performance computer cluster (under project Nos n9244kk), owned by the University of Oslo and the Norwegian metacenter for High Performance Computing (NOTUR), and operated by the Department for Research Computing at USIT, the University of Oslo IT-department. The project was supported by the Research Council of Norway (grant number 222378/F20 to KSJ).

## Author contribution

SM, KSJ and SJ designed the study and obtained financial support; XJ, BM, VM and SM participated in the experimental design, macrophage cell culture development and sampling; XJ, BM, OKT and MHS carried out bioinformatic analyses and posterior statistical analysis. All authors participated in drafting the final manuscript.

## Data availability

RNA-Seq raw data generated in this study have been deposited in the NCBI Sequence Read Archive (SRA) under BioProject accession code PRJNA299100 (Bac and dsRNA done, reads for MDP and IsoSeq is uploading)

## Supporting Information

Supplementary Figure.1. Phagocytosis assays on macrophage cell cultures from 4 individuals.

Supplementary Figure.2. Extended results for RNA-Seq of cod macrophages post immune challenge by dsRNA and bacterial PAMPs (Bac).

Supplementary Figure.3. Extended results for RNA-Seq of cod macrophages post MDP time-course activation.

Supplementary Figure.4. Overlapping analysis of DEGs activated by different MAMPs. Supplementary Figure.5. Biological role of dsRNA and MDP-responsive genes Supplementary Figure 6. Gene tracks of representative PRRs of Atlantic cod in response to dsRNA and MDP challenges.

Supplementary Table.S1. Phagocytosis results for both untreated and microbial co-cultured cod macrophages.

Supplementary Table.S2 Overlapping analysis of MAMP-dependent DEGs with annotated human gene identifiers (gene_names) via Venn diagram for Figs. 2 *C*, 5 and S4

Supplementary Table.S3 Species and sequences information for synteny Supplementary Table.S4. Literatures for functional studies of RLRs-mediated antiviral response in indicated vertebrates

Supplementary Table.S5 Summary of statistics

Supplementary Table.S6 Dosage of different MAMPs and inhibitors used in cod macrophages immune challenge.

Supplementary Table.S7 cDNA libraries for screening of cod macrophages response to MAMP challenges.

Supplementary Table.S8 Oligo nucleotides primer sequences for target genes

Supplementary Table.S9 Accuracy estimation of transcriptomic assemblies of Cufflinks and StringTie compared with cod genome reference annotation “gadMor2” by Gffcompare.

## Supplementary Data

**Supplementary Data 1 |** Differential expression analysis and GOs/KEGGs enrichment output

