## Supplementary figures and images for "Innovation in NLR and TLR sensing drives the MHC-II free Atlantic cod immune system"

### Figure S1

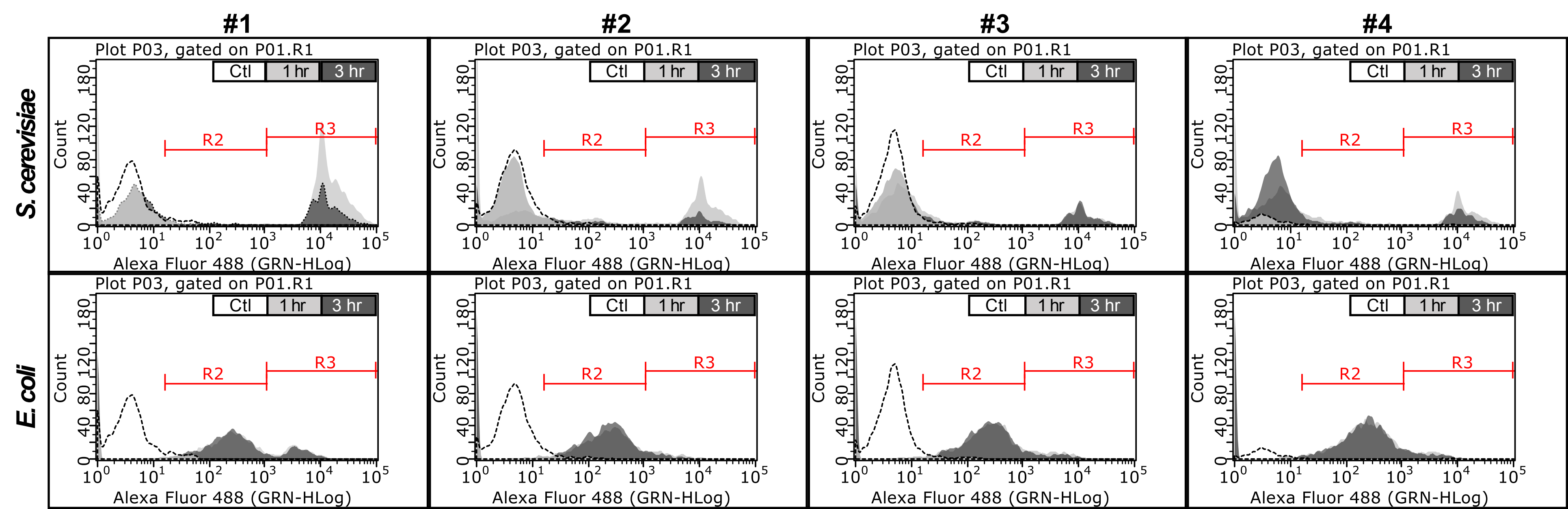

### Figure S2

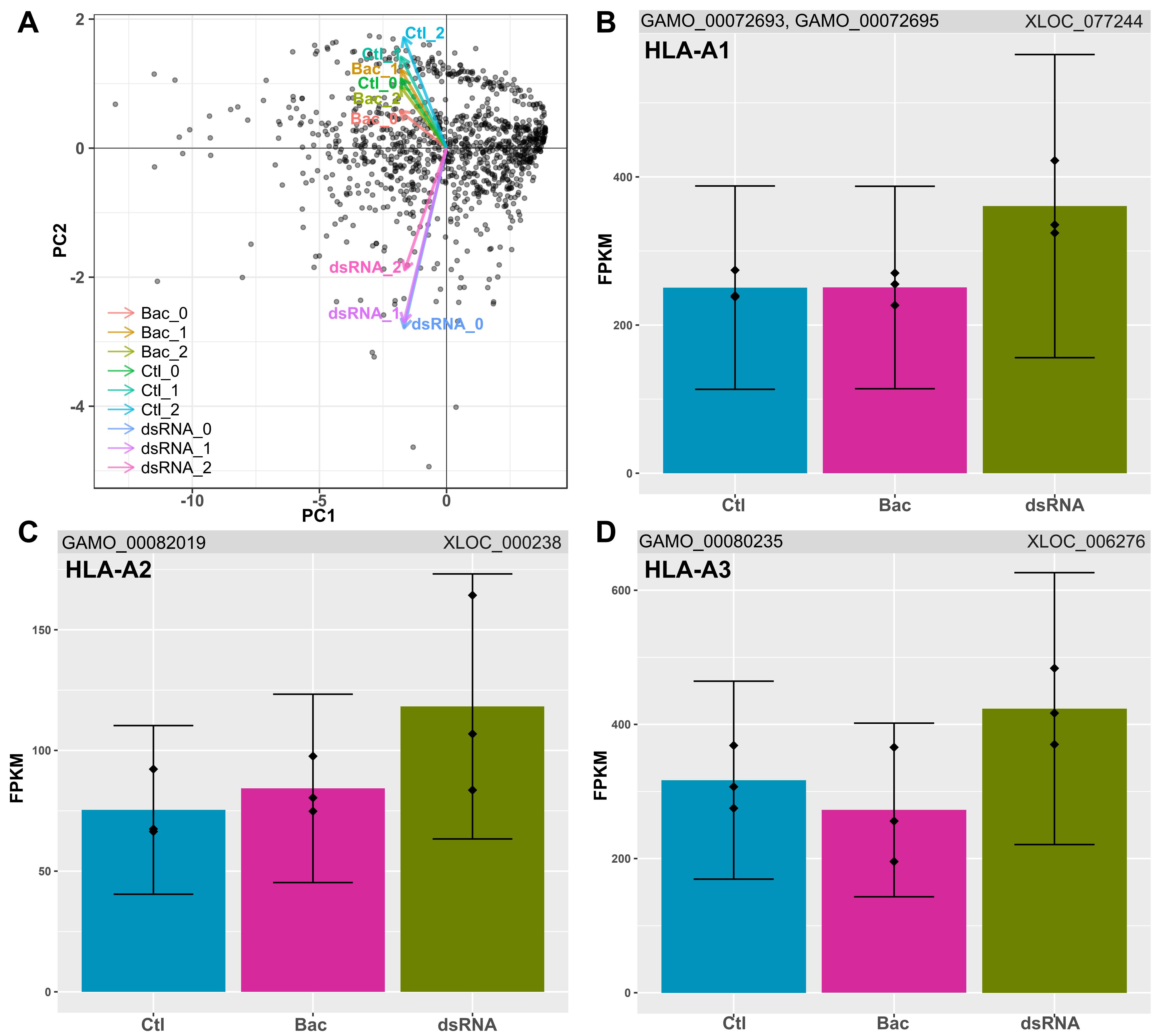

### Figure S3

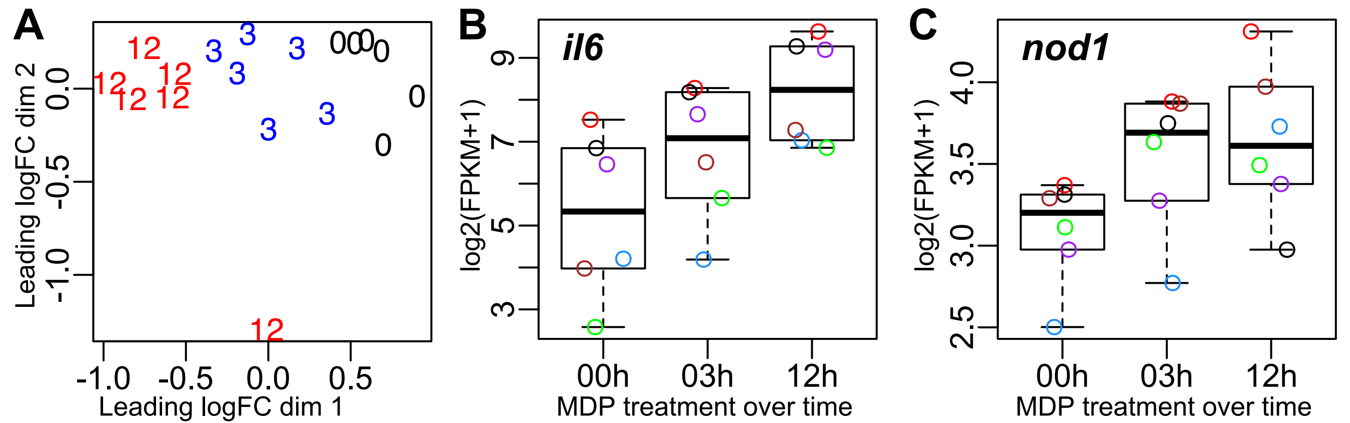

### Figure S4

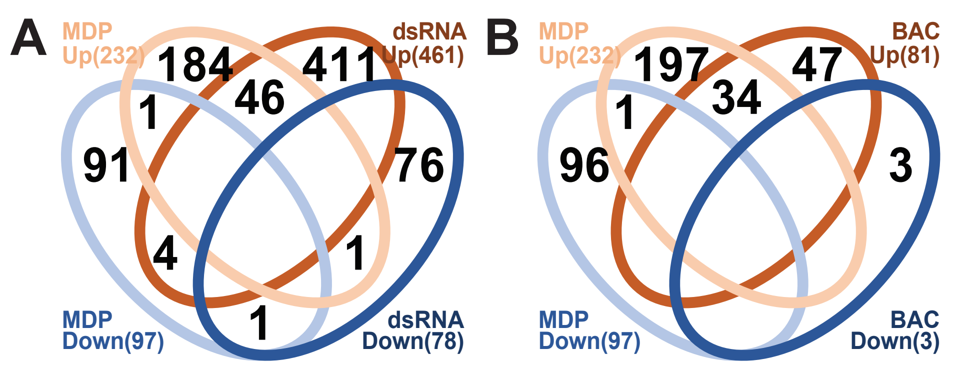

### Figure S5

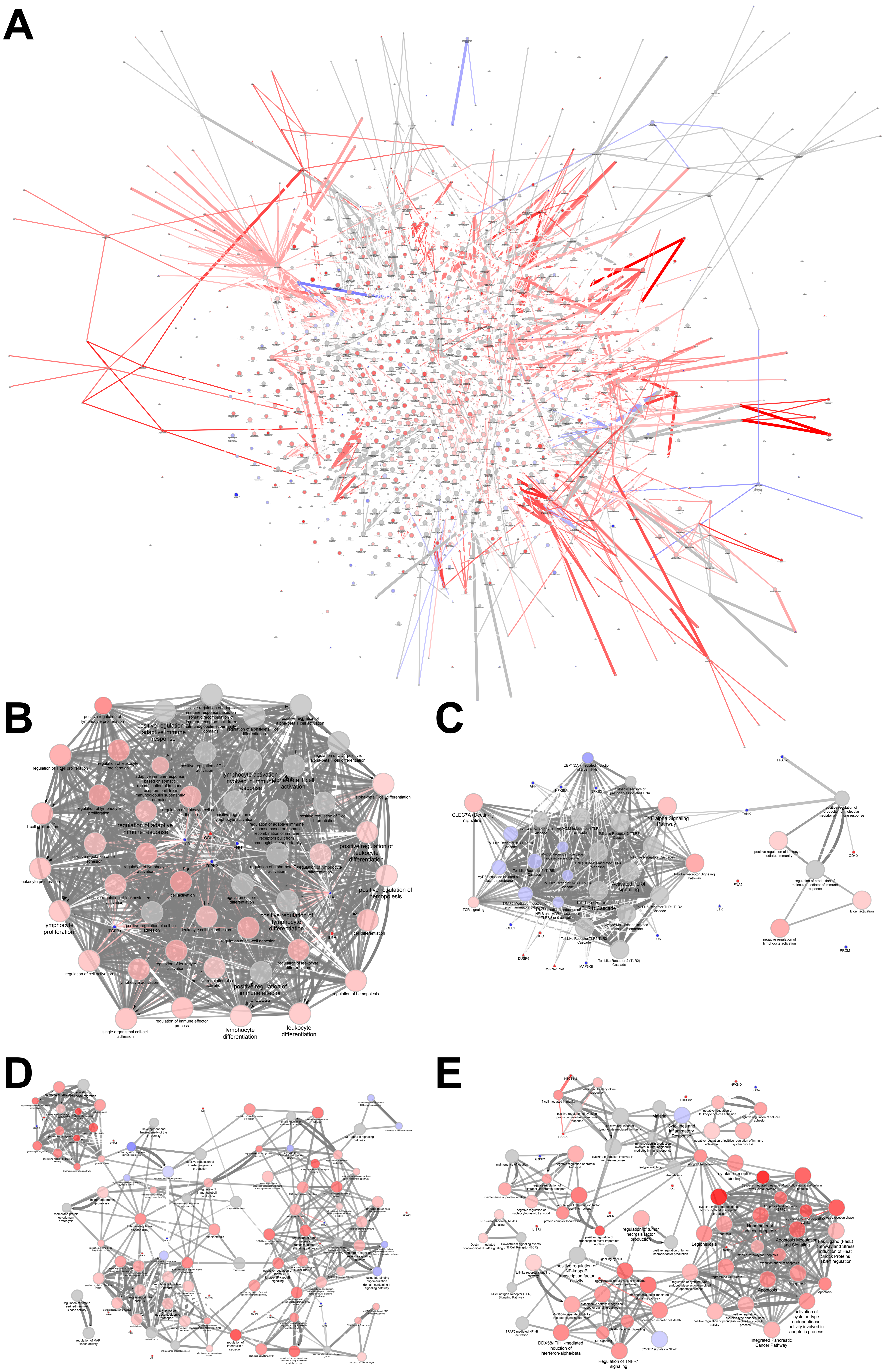

### Figure S6

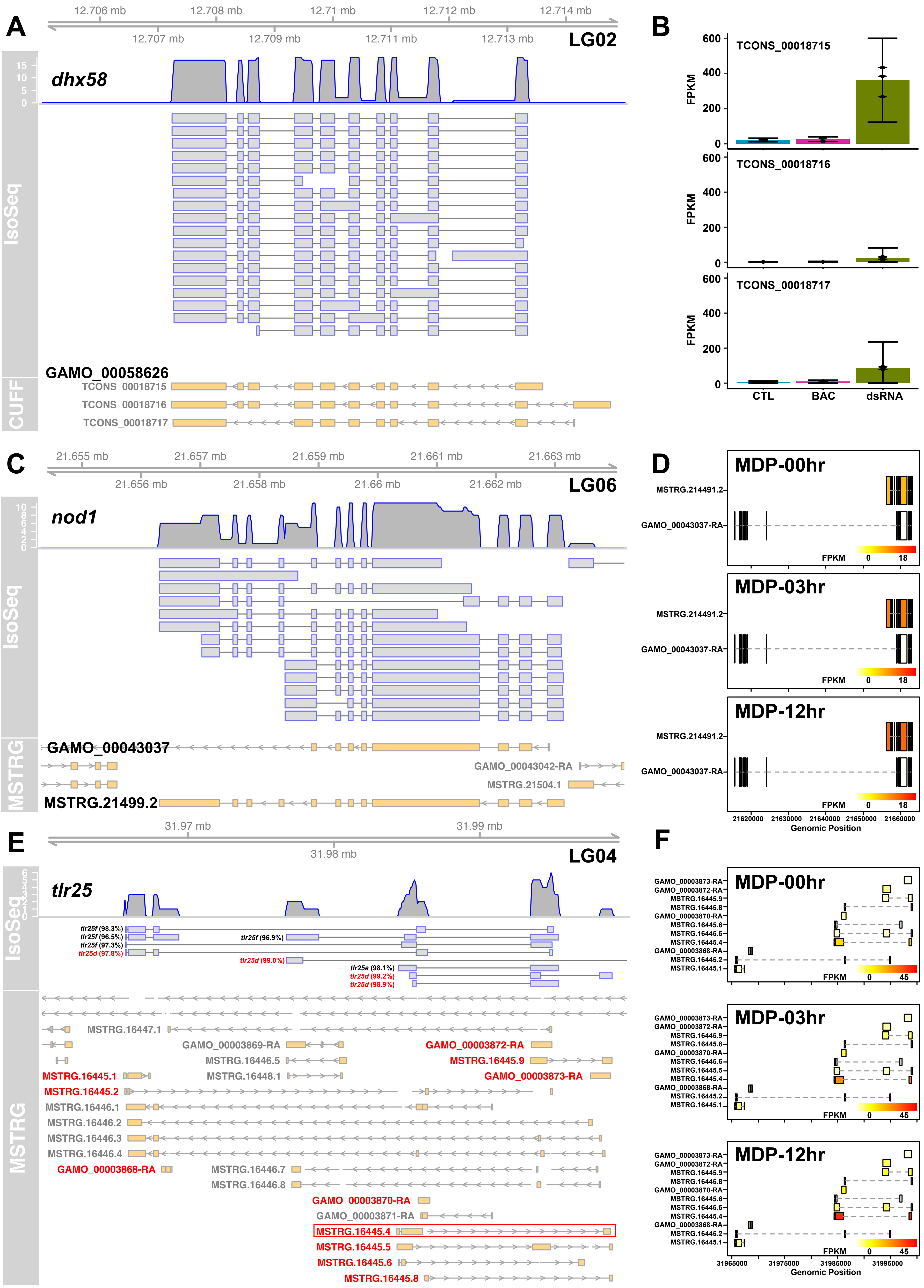

### Figure S7

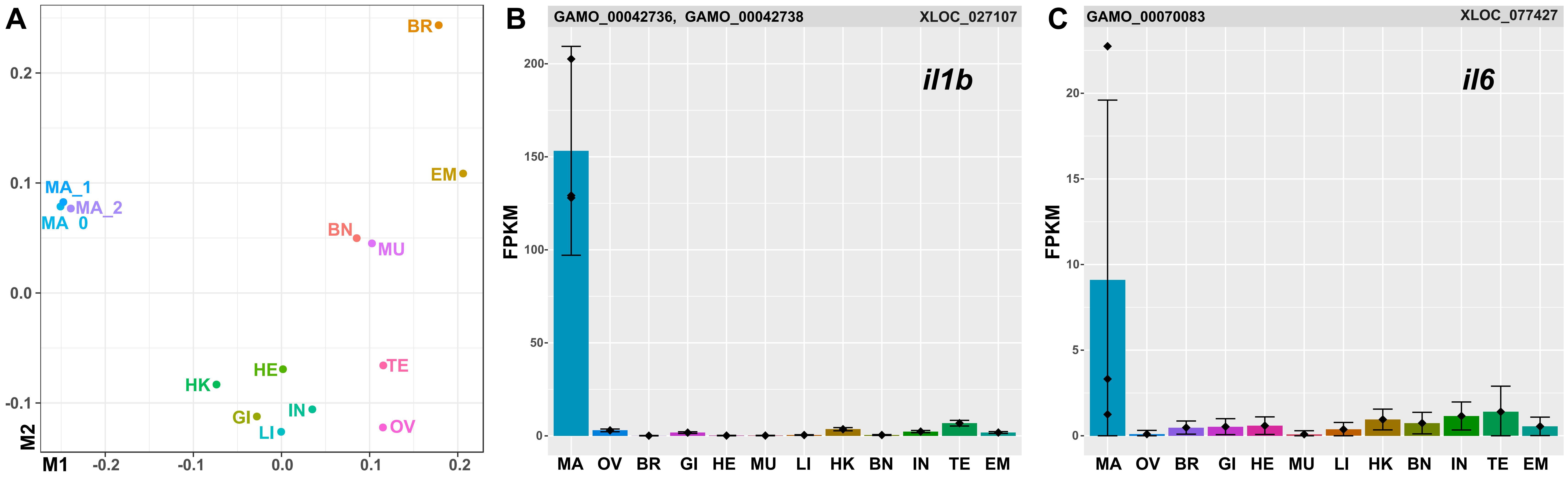
